# Detection of Candidate Circular RNAs to Monitor Anti-Hormonal Response in the Mammary Gland

**DOI:** 10.64898/2026.03.26.714379

**Authors:** Nico Trummer, Malte Weyrich, Paige Ryan, Priscilla A. Furth, Markus Hoffmann, Markus List

## Abstract

Anti-hormonal therapies such as selective estrogen receptor modulators like tamoxifen or aromatase inhibitors like letrozole represent a cornerstone for breast cancer prevention and therapy of estrogen receptor-positive breast cancer. Therapeutic monitoring can include blood tests and imaging; however, genetically-based approaches are not yet in practice. Ideally, a test would be able to detect a positive molecular response across different estrogen pathway-suppressive approaches. Circular RNAs are a species of non-coding RNAs detectable in plasma that have been proposed as non-invasive therapeutic biomarkers. To determine whether a set of specific circular RNAs is altered across estrogen-suppressive pathway approaches, we analyzed mammary gland-specific total RNA sequencing data from two individual genetically engineered mouse models (GEMMs) of estrogen pathway-induced breast cancer, with or without exposure to tamoxifen or letrozole. The nf-core/circrna pipeline was used to identify circRNAs that were differentially expressed in response to either tamoxifen or letrozole. We then screened for circRNAs that were differentially regulated by both anti-hormonals. Four up-regulated and 31 down-regulated circRNAs with host genes known to be expressed in human breast epithelial cells were identified as showing reproducible differential regulation in response to anti-hormonal treatment.

## Introduction

Circular RNAs (circRNAs) represent a species of generally non-coding RNAs that have the capacity to serve as biomarkers, therapeutic targets and to influence downstream gene expression networks (*1*, *2*). CircRNAs are primarily classified as long non-coding RNAs (lncRNAs), although a few of them encode proteins (*3*). They are formed through a back-splicing event of precursor messenger RNA (pre-mRNA), resulting in the fusion of 5’ and 3’ ends of a portion of pre-mRNA that is spliced out (*4*). Their closed circular loop structure makes them less prone to degradation (*5*). circRNAs can be transported to the extracellular space in exosomes (*6*, *7*). Eventually, tissue-generated circRNAs that are secreted in exosomes can be found circulating in the blood and are considered possible biomarkers of response that have the benefit of being assessed using liquid biopsy (*8*). CircRNAs are generally expressed at levels between two and 10% of that of their linearly expressed host genes, and it is possible for a single host gene to use alternative back-splicing mechanisms to produce multiple circRNA isoforms (*6*, *7*). In some circumstances, multiple circRNAs share the same back-splice junction (*9*). For use as biomarkers, previous studies have documented that a minimum of a statistically significant 2-fold change (log2fold change of one) is recommended, with higher fold changes proving more reliable, both up- and down-regulated biomarkers are useful, and the use of three or more individual circRNA biomarkers in combination provides a more robust analysis (*10*, *11*)

Anti-hormonal selective estrogen modulators (SERMs) such as tamoxifen and aromatase inhibitors (AIs) such as letrozole represent two of the major therapeutic classes of drugs used for the reduction of breast cancer risk and/or therapy of estrogen receptor-positive breast cancer (*12*). Here, we investigated the hypothesis that there would be common circRNAs that would be differentially regulated by both classes of antihormonals, which would represent candidate circRNAs that could be monitored for a positive response to anti-hormonal therapy. The nf-core/circrna detection pipeline (*13*, *14*) was used to identify genetic regions containing BSJs representing candidate circRNAs that were differentially regulated following anti-hormonal exposure. The datasets analyzed were total RNA sequencing data of mammary gland tissue derived from a time-course experiment that employed two genetically engineered mouse models (GEMM) of estrogen pathway-induced breast cancer that exhibited the expected loss of mammary gland cellularity and decrease in preneoplasia and neoplasia following exposure to either tamoxifen or letrozole (*15*). Previous published work has concentrated on identifying circRNAs whose expression is associated with tamoxifen resistance (*16–19*), and, in the case of AIs, circRNAs that are predictive of a durable AI response (*20*). Here, we extend our understanding of the relationship between circRNAs and anti-hormonals by identifying candidate circRNAs that are differentially regulated in the context of a positive therapeutic response in whole mammary tissue. Mammary gland tissue from GEMM have been shown to be informative preclinical molecular models of genetic response to therapeutic interventions for human breast cancer prevention and treatment (*21*, *22*). CircRNAs can have orthologous structures in different species, such as mice and humans. These orthologous circRNAs may exhibit conserved genomic origins or have emerged convergently, with variation in conserved function and expression (*23*, *24*).

Here, we used a bioinformatic approach (see Materials and Methods for details) to identify four upregulated and 31 downregulated regions containing BSJs representing candidate circRNAs commonly differentially regulated by both tamoxifen and letrozole in the mouse mammary gland. Two of the up-regulated and 24 of the down-regulated candidate circRNAs have analogous regions in humans containing previously identified circRNAs, representing a first set of potential circRNA biomarkers that could be used to assess anti-hormonal response in human breast tissue at risk for breast cancer development. Five additional candidate circRNA regions identified (one up-regulated and four down-regulated) that have host genes known to be expressed in human breast cells but are not currently listed in any human circRNA database may represent newly identified circRNA structures responsive to antihormonal therapy.

## Results

### Detection of differentially expressed candidate circRNA regions in mouse mammary gland following exposure to tamoxifen or letrozole

For the detection of candidate tamoxifen-regulated circRNAs, total RNAseq data from mammary glands of both mammary tumor virus–reverse tetracycline–controlled transactivator (*MMTV-rtTA*)/Tet-operator (*tet-op*)–*Esr1* and mouse mammary tumor virus–reverse tetracycline–controlled transactivator/*tet-op*–*CYP19A1* mice with and without tamoxifen exposure were compared. For the detection of candidate letrozole regulated circRNAs, total RNAseq data from mammary glands of both *MMTV-rtTA/tet-op-Esr1* and *MMTV-rtTA/tet-op-CYP19A1* with and without letrozole exposure were compared. The two different GEMMs analyzed were chosen to represent two major, distinctly different molecular pathways underlying estrogen pathway-related human breast cancer generation, Estrogen Receptor alpha over-expression as compared to aromatase over-expression that increases mammary gland localized estrogen levels (*15*, *25–27*). Application of the nf-core/circrna detection pipeline in combination with DESeq2 resulted in the identification of 61 candidate circRNA regions differentially regulated by tamoxifen exposure and 51 candidate circRNAs differentially regulated by letrozole exposure (adj. p≤0.05) (Figure 1). Data were examined to determine how many of these candidate circRNAs were differentially expressed in the same direction (either up-regulated or down-regulated) by both tamoxifen and letrozole. Thirty-five circRNAs commonly differentially regulated by both tamoxifen and letrozole were identified (Figure 2). Chromosomal coordinates of the candidate circRNA regions, names of candidate host genes within the chromosomal coordinates were determined, and annotation to murine circRNA databases (CircAtlas, CircBase, CircBank, Circpedia, Tissue-Specific CircRNA Database (TSCD) was performed (Tables 1-4). Overall, 33 of the 35 detected circRNAs (94%) in mouse mammary tissue were supported by entries in at least one of the murine circRNA databases, indicating that the pipeline was able to detect true circRNAs as defined by their record in established databases.

**Figure 1.**
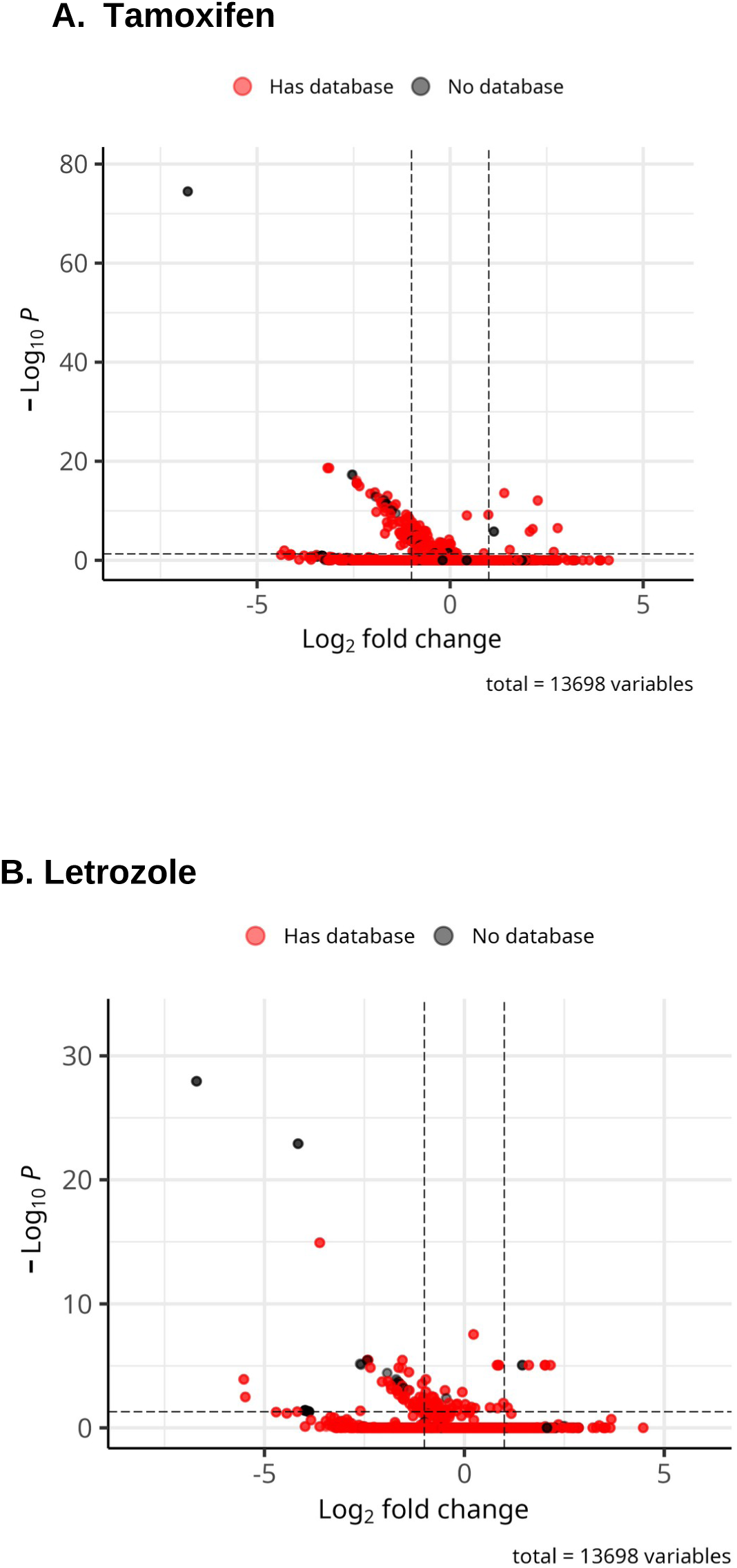
Volcano plots of significantly expressed circRNAs after pre-filtering and lfc shrinkage for both letrozole and tamoxifen. **A.** Volcano plot of significantly expressed circRNAs following tamoxifen exposure. **B.** Volcano plot of significantly expressed circRNAs following letrozole exposure. Any circRNA having a padj < 0.05 and abs(lfc) > 1 was considered significant. CircRNAs that are highlighted red indicate there is an annotation in either circAtlas (*53*) circBase (*52*), or the lifted circBase database as described in the methods section.

**Figure 2.**
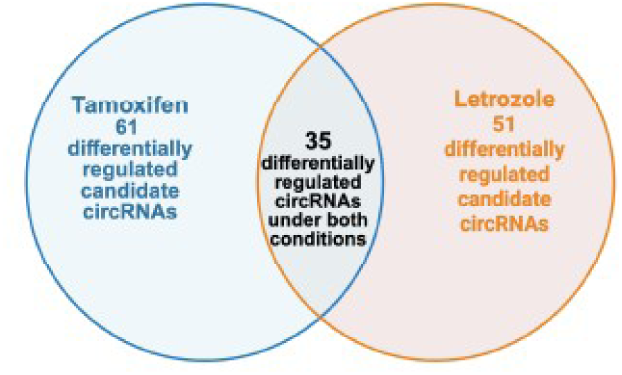
Identification of candidate circRNAs significantly differentially expressed with both tamoxifen and letrozole exposure. Venn diagram of the overlap of differentially expressed circRNAs with tamoxifen versus letrozole exposure. Twenty-one differentially expressed candidate circRNAs were identified using the nf-core/circRNA pipeline and Dseq2 to identify differentially expressed circRNAs under different hormonal conditions (padj. <0.05). Created in BioRender: Furth, P. (2026) https://BioRender.com/x0xbrcl

### Characterization of candidate circRNA regions commonly differentially regulated by exposure to tamoxifen and letrozole

Four candidate circRNA regions were found to be significantly up-regulated and 31 candidate circRNA regions were found to be significantly down-regulated and by exposure to both tamoxifen and letrozole (padj<0.05) (Tables 1, 2). Predicted host genes derived from the chromosomal coordinates of the candidate circRNA regions, basemean (average normalized count of the candidate region across all samples), log2foldchange (measure of how much a value has changed between two conditions expressed on a logarithmic scale), lfcSE (standard error estimate for the log2 fold change estimate), stat (log2FoldChange divided by lfcSE, which is compared to a standard normal distribution to generate a two-tailed pvalue), pvalue (probability of observing the measured difference in gene expression between groups assuming the null hypothesis, uses the Wald test), and padj (adjusted p-value) were determined. Average normalized count values (base mean) ranged from 0.46 to 1.58 for up-regulated and 1.03 to 214.05 for down-regulated candidate circRNAs with statistically significant log2fold changes ranging from 1.13 to 2.78 for up-regulated and −1.02 to −6.79 for down-regulated candidate circRNAs (Tables 1, 2). Two candidate down-regulated circRNAs mapping to a transcript set (Rn18s-rs5, Gm26917, AY036118) were marked individually by the pipeline but differed by only two nucleotides at the start and therefore appeared to represent the same general circRNA structure, leaving a total of 30 individual candidate down-regulated circRNAs. In two cases, the candidate circRNA regions mapped to two adjacent genes. One was an up-regulated region on chromosome 7 mapped to *Gm34744* (a predicted long non-coding (lnc) RNA in mice) and *Bcam. Gm34744* is. One down-regulated region on chromosome 6 mapped to *Gm52873* (predicted mouse gene) and *Washc2*. A relative comparison of baseline host gene in transcripts per million (TPM) and basemean circRNA expression levels showed Circular-to-Linear Ratios (CLR) ranging from 0.002 to 0.60 (Supplementary Tables 1, 2). All of the identified circRNAs revealed a fold-change of greater than 2-fold, and 4-fold changes were documented for 20 of the 31 down-regulated circRNAs (Table 2), meaning that the collection of circRNAs identified included multiple candidates that could be used for a combination circRNA biomarker platform. Host gene expression was examined to determine if the differentially regulated circRNA expression identified might be secondary to significant changes in host gene expression following exposure to tamoxifen or letrozole. No instance of host gene expression values paralleling circRNA expression levels across both anti-hormonal exposures and both models was identified although it was noted that for the up-regulated circRNA from host gene *Bcam,* mammary glands from *MMTV-rtTA/tet-op-Esr1* mice showed significantly increased host gene expression with both tamoxifen and letrozole, although the changes were less than 2-fold (Supplementary Table 1). Similarly, the down-regulated circRNAs mapped to host genes *Rpn1* and *Strbp* showed significant decreases with both tamoxifen and letrozole in the mammary glands from *MMTV-rtTA/tet-op-Esr1* mice, but the changes were less than 2-fold, while in all cases the changes in the circRNA expression levels were greater than 2-fold (Supplementary Table 2). Finally, the host gene set belonging to the circRNAs identified was then assessed through Gene Set Enrichment Analysis (GSEA) through both mouse and human gene sets to determine if the Molecular Signatures Database (MSigDB) would identify the set as having any statistically significant relationships to established gene sets. Statistically significant relationships to C3 (*28*) regulatory target gene sets, C2 curated gene sets, and M3 regulatory target gene sets were identified (data not shown) but not for any hallmark gene sets particularly either HALLMARK_ESTROGEN_RESPONSE_EARLY(M5907) or HALLMARK_ESTROGEN_RESPONSE_LATE(M5906) gene sets. It is known that circRNAs can show differential expression independent of host gene expression patterns (*29*, *30*). Not only are the majority (94%) of identified differentially regulated circRNAs in one of the established mouse circRNA databases but those that are listed are all present in at least two of the queried databases and 12 of the 33 identified circRNAs mapped to more than one listing in at least one of the databases examined (Tables 3, 4). To determine which of the candidate circRNAs identified in mouse mammary gland tissue were plausible candidate circRNA biomarkers for human breast tissue, host gene expression in human breast tissue was examined and identification of orthologous human circRNAs in human circRNA databases was performed. Host genes for three of the four up-regulated and 27 of the 30 down-regulated circRNAs (overall 88%) were documented to be expressed in human breast primary epithelial cells (Tables 5, 6). Human orthologous circRNAs were identified for 25 of the 30 host genes expressed in human breast. Five of the orthologous circRNAs have been previously documented as being expressed in human breast (n=4 in CircAtlas: hsa-FMN1_0002, hsa-CSNK1G3_0001, hsa-ZNF148_0018, hsa-MBNL1 and n=1 in TCSD: circRpn1).

**Table 1.**
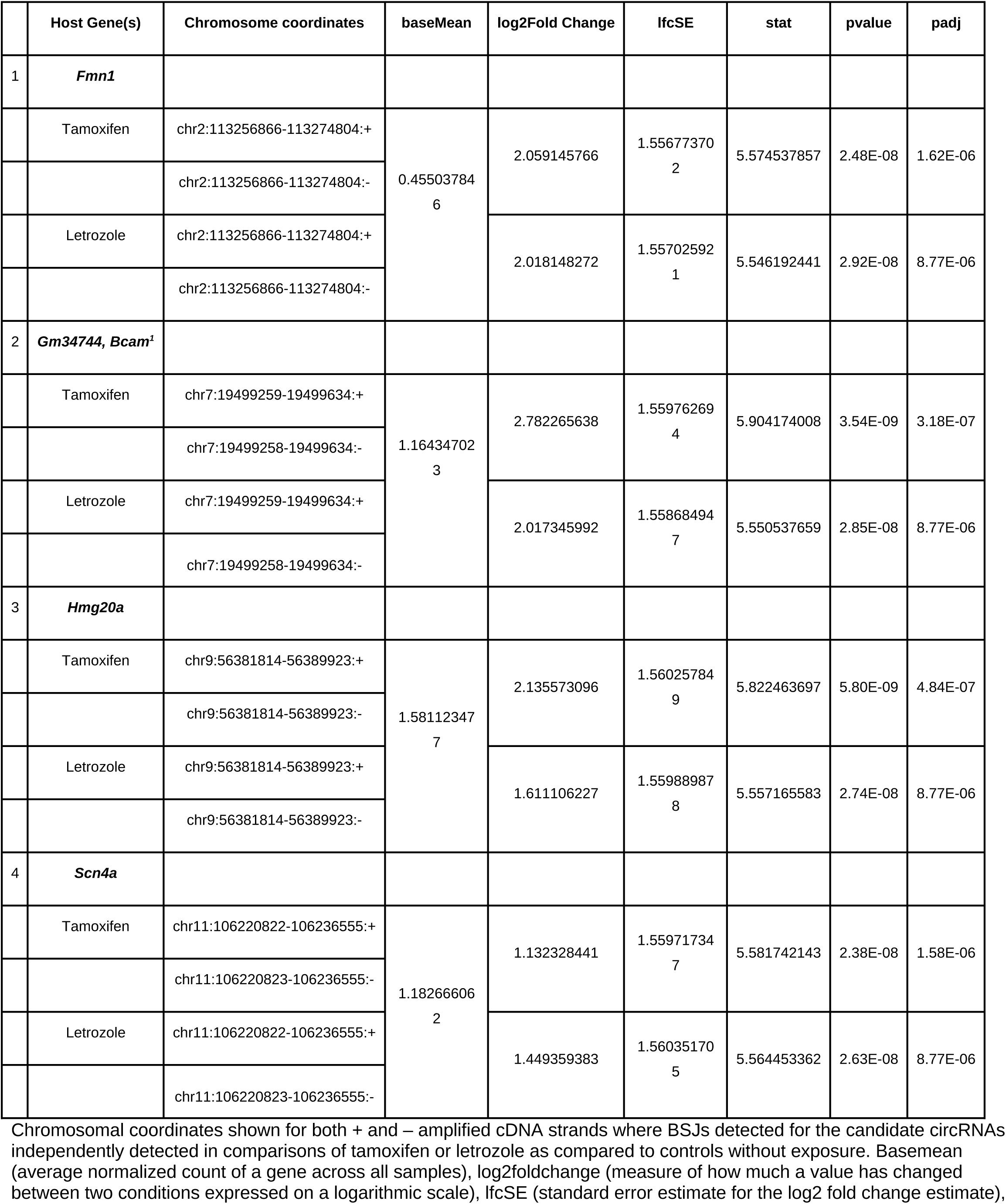
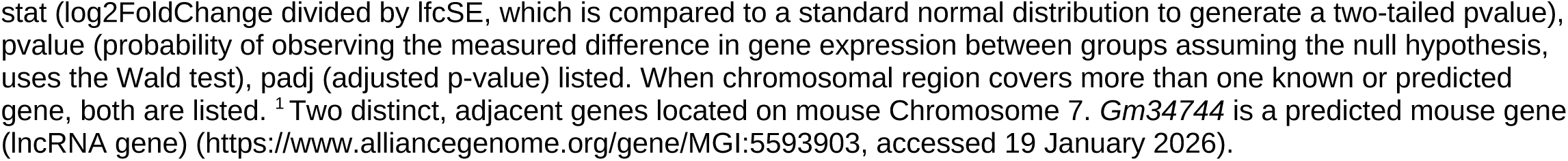
Candidate circRNA regions identified as commonly up-regulated by tamoxifen and letrozole exposure in mouse mammary gland.

|  | Host Gene(s) | Chromosome coordinates | baseMean | log2Fold Change | lfcSE | stat | pvalue | padj |
| --- | --- | --- | --- | --- | --- | --- | --- | --- |
| 1 | ***Fmn1*** |  |  |  |  |  |  |  |
|  | Tamoxifen | chr2:113256866-113274804:+ | 0.45503784<br>6 | 2.059145766 | 1.55677370<br>2 | 5.574537857 | 2.48E-08 | 1.62E-06 |
|  |  | chr2:113256866-113274804:- |  |  |  |  |  |  |
|  | Letrozole | chr2:113256866-113274804:+ |  | 2.018148272 | 1.55702592<br>1 | 5.546192441 | 2.92E-08 | 8.77E-06 |
|  |  | chr2:113256866-113274804:- |  |  |  |  |  |  |
| 2 | ***Gm34744, Bcam<sup>1</sup>*** |  |  |  |  |  |  |  |
|  | Tamoxifen | chr7:19499259-19499634:+ | 1.16434702<br>3 | 2.782265638 | 1.55976269<br>4 | 5.904174008 | 3.54E-09 | 3.18E-07 |
|  |  | chr7:19499258-19499634:- |  |  |  |  |  |  |
|  | Letrozole | chr7:19499259-19499634:+ |  | 2.017345992 | 1.55868494<br>7 | 5.550537659 | 2.85E-08 | 8.77E-06 |
|  |  | chr7:19499258-19499634:- |  |  |  |  |  |  |
| 3 | ***Hmg20a*** |  |  |  |  |  |  |  |
|  | Tamoxifen | chr9:56381814-56389923:+ | 1.58112347<br>7 | 2.135573096 | 1.56025784<br>9 | 5.822463697 | 5.80E-09 | 4.84E-07 |
|  |  | chr9:56381814-56389923:- |  |  |  |  |  |  |
|  | Letrozole | chr9:56381814-56389923:+ |  | 1.611106227 | 1.55988987<br>8 | 5.557165583 | 2.74E-08 | 8.77E-06 |
|  |  | chr9:56381814-56389923:- |  |  |  |  |  |  |
| 4 | ***Scn4a*** |  |  |  |  |  |  |  |
|  | Tamoxifen | chr11:106220822-106236555:+ | 1.18266606<br>2 | 1.132328441 | 1.55971734<br>7 | 5.581742143 | 2.38E-08 | 1.58E-06 |
|  |  | chr11:106220823-106236555:- |  |  |  |  |  |  |
|  | Letrozole | chr11:106220822-106236555:+ |  | 1.449359383 | 1.56035170<br>5 | 5.564453362 | 2.63E-08 | 8.77E-06 |
|  |  | chr11:106220823-106236555:- |  |  |  |  |  |  |

Chromosomal coordinates shown for both + and – amplified cDNA strands where BSJs detected for the candidate circRNAs independently detected in comparisons of tamoxifen or letrozole as compared to controls without exposure. Basemean (average normalized count of a gene across all samples), log2foldchange (measure of how much a value has changed between two conditions expressed on a logarithmic scale), lfcSE (standard error estimate for the log2 fold change estimate),
stat (log2FoldChange divided by lfcSE, which is compared to a standard normal distribution to generate a two-tailed pvalue), pvalue (probability of observing the measured difference in gene expression between groups assuming the null hypothesis, uses the Wald test), padj (adjusted p-value) listed. When chromosomal region covers more than one known or predicted gene, both are listed. <sup>1</sup>Two distinct, adjacent genes located on mouse Chromosome 7. *Gm34744* is a predicted mouse gene (lncRNA gene) (https://www.alliancegenome.org/gene/MGI:5593903, accessed 19 January 2026).

**Table 2.**
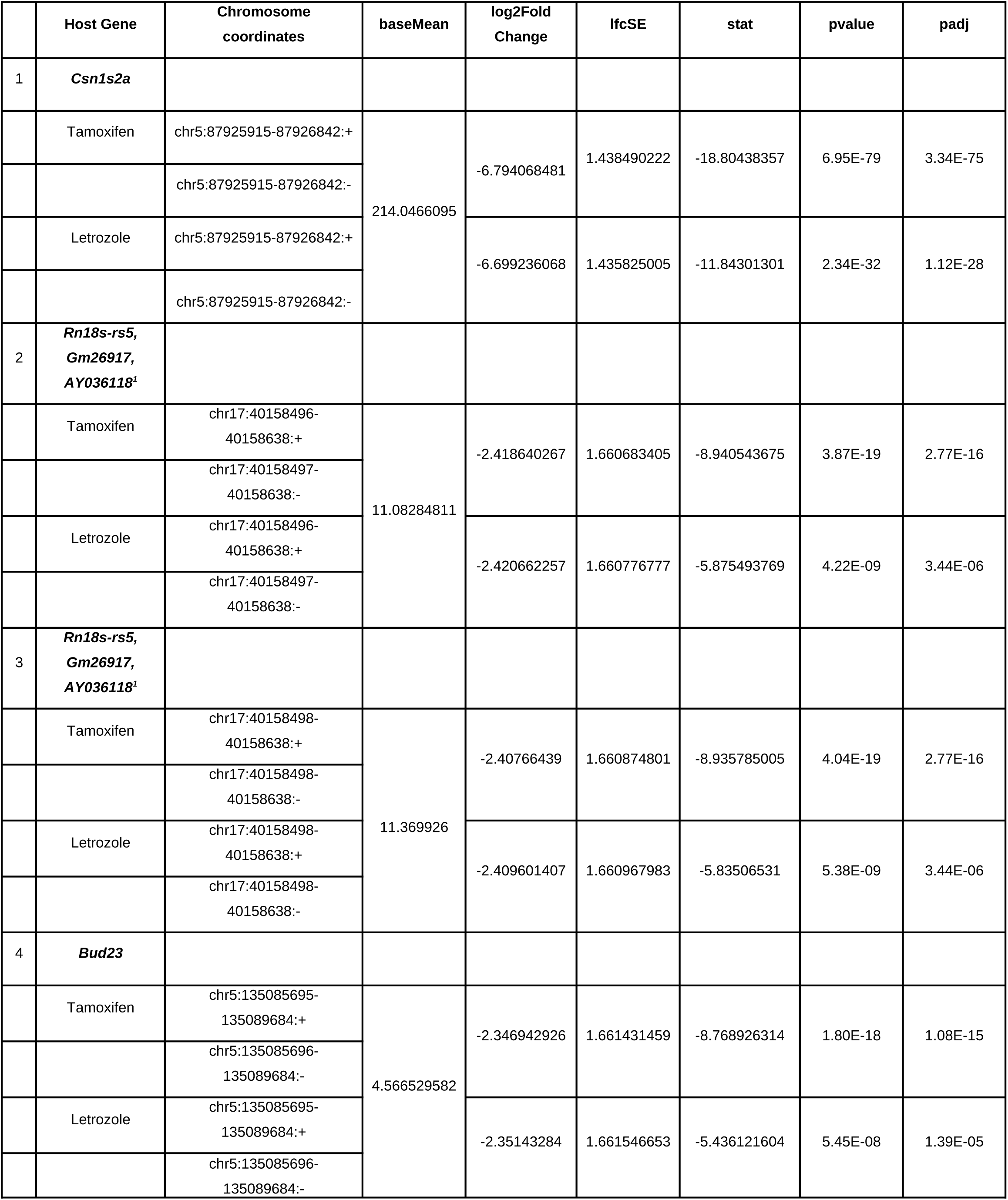
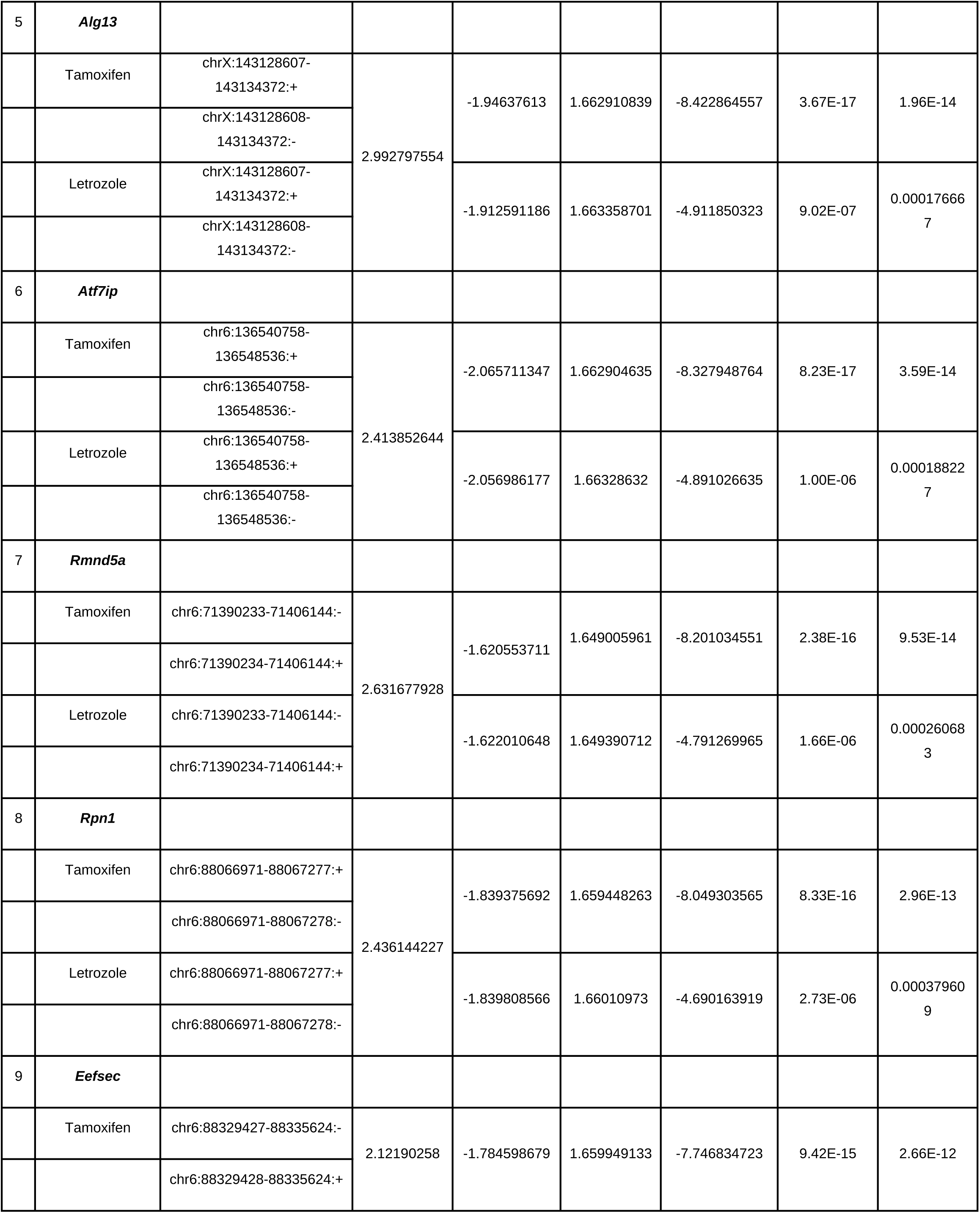
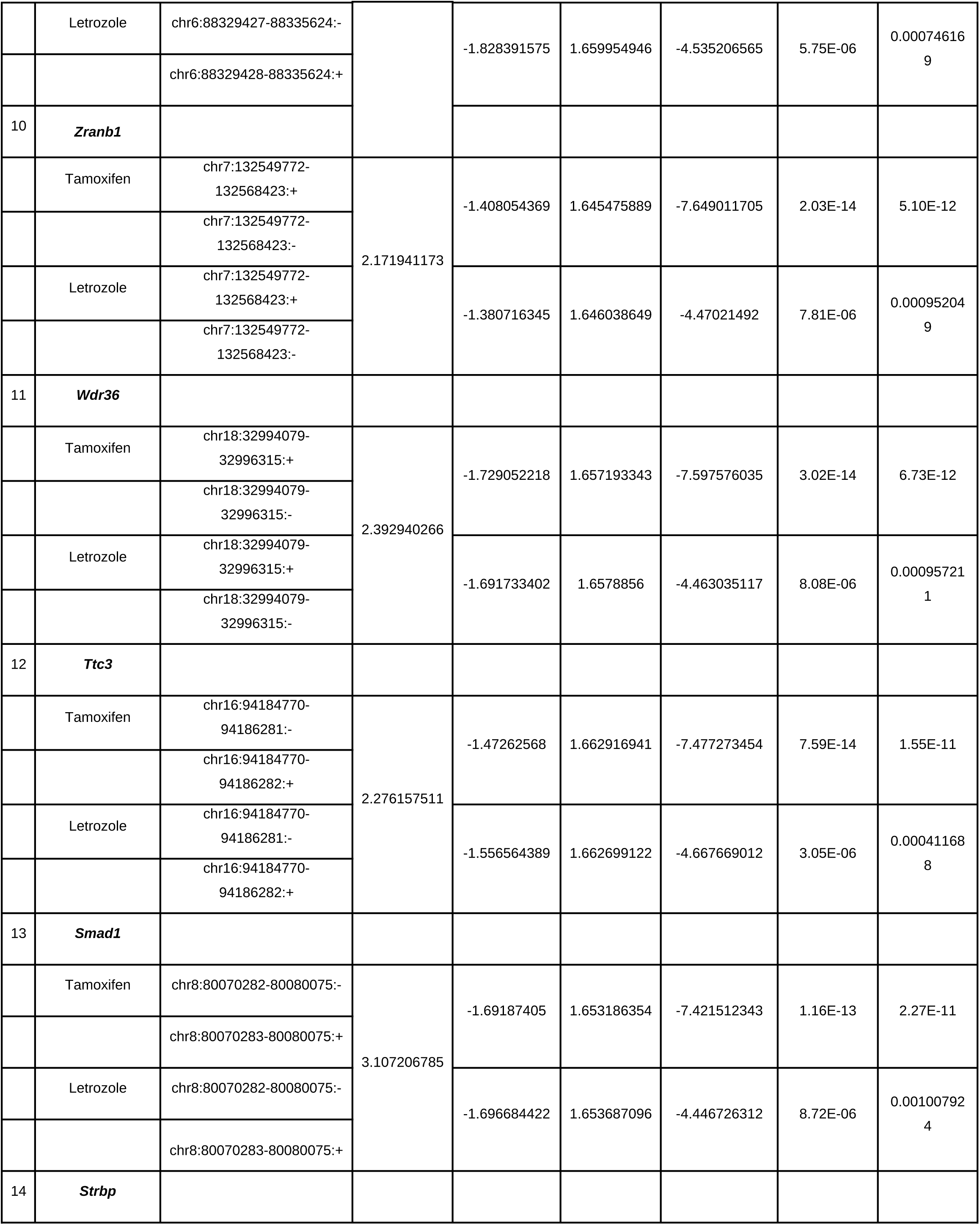
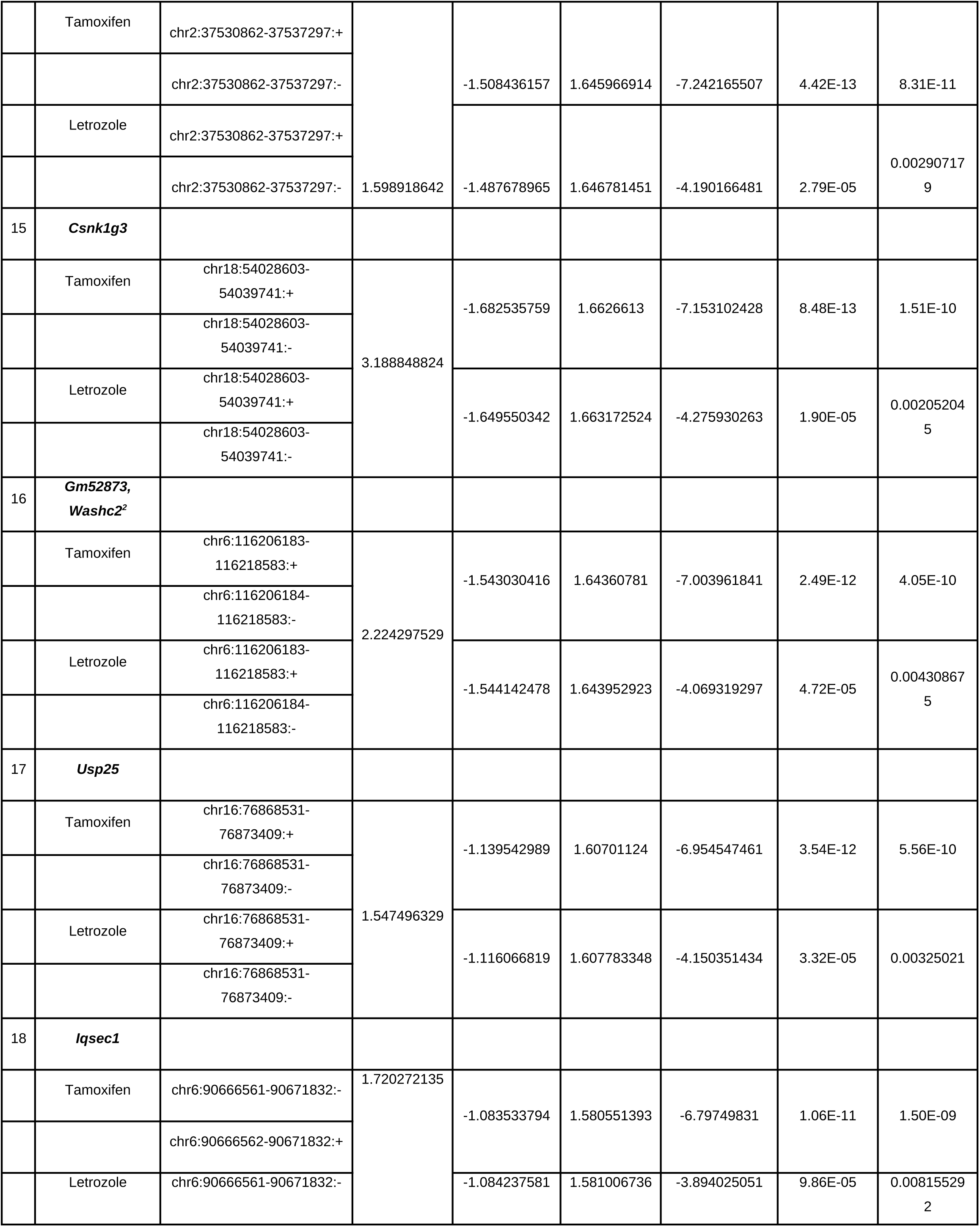
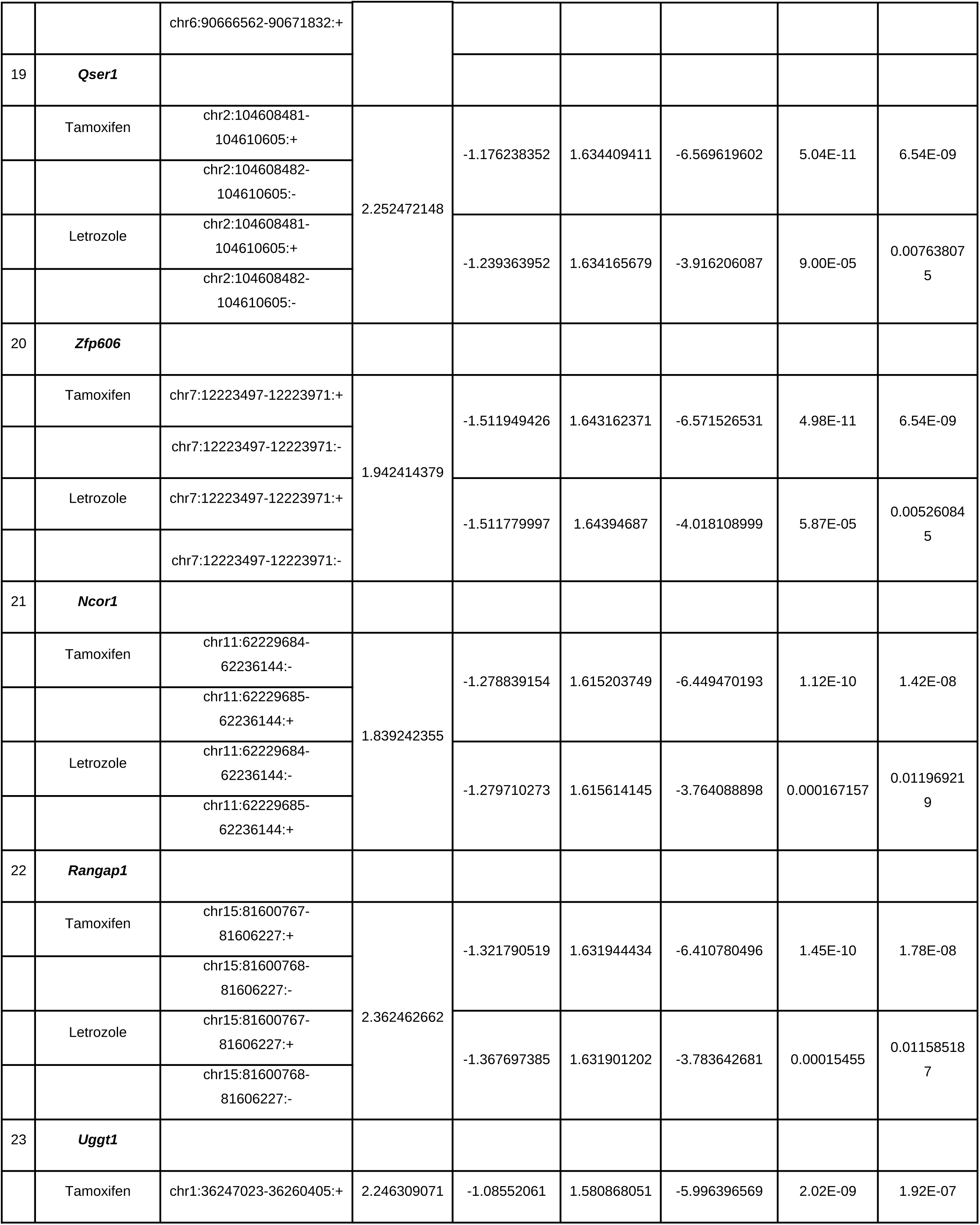
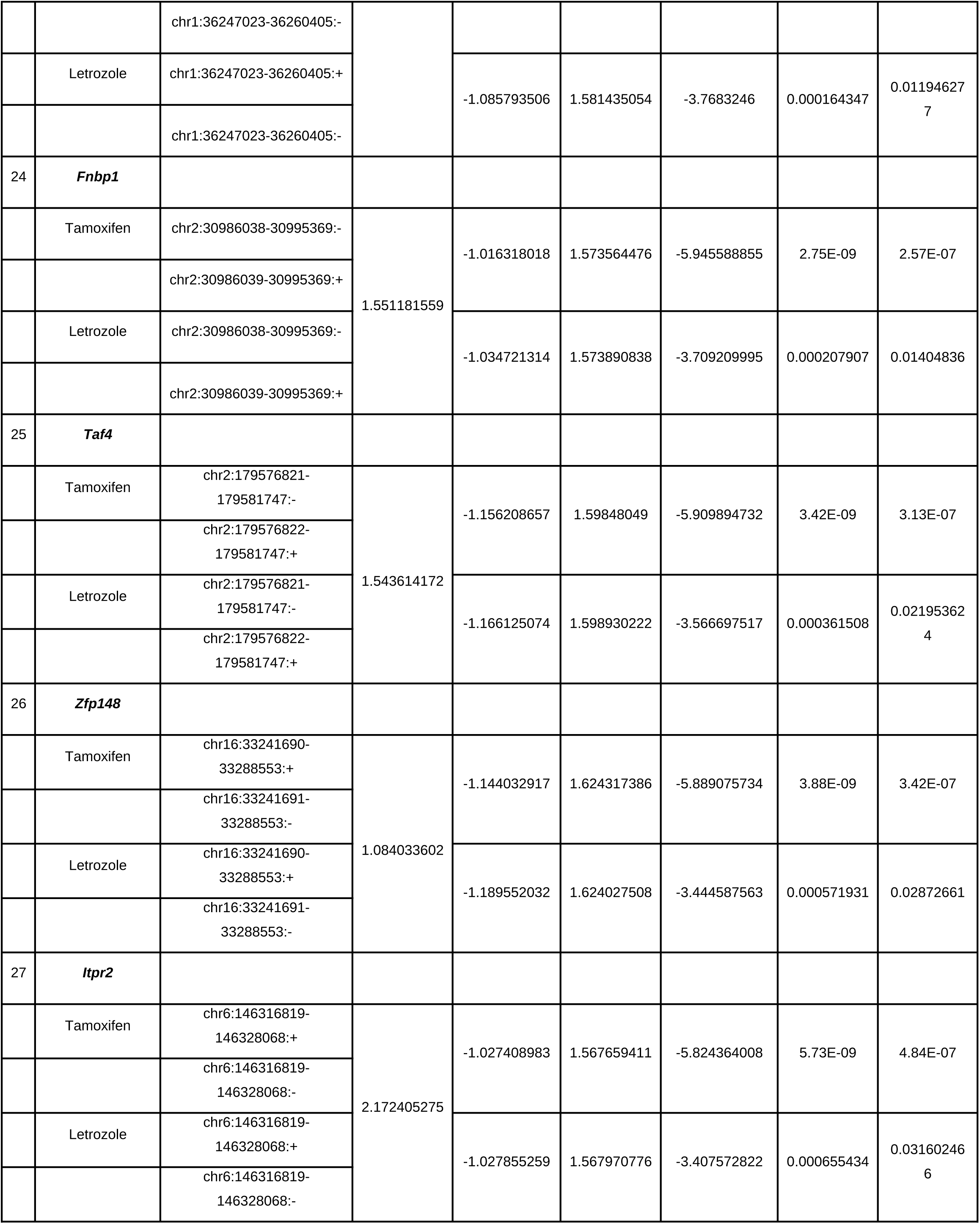
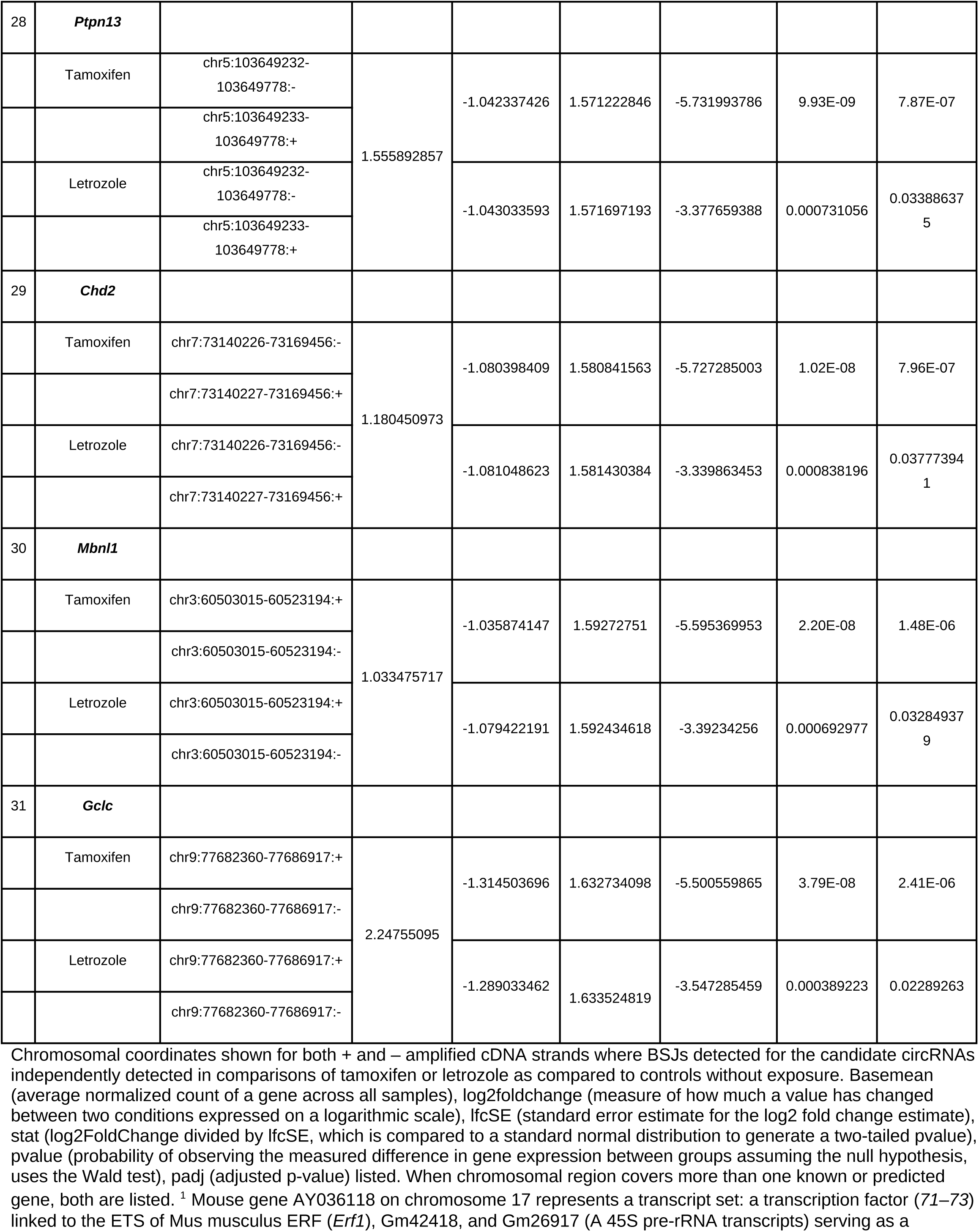
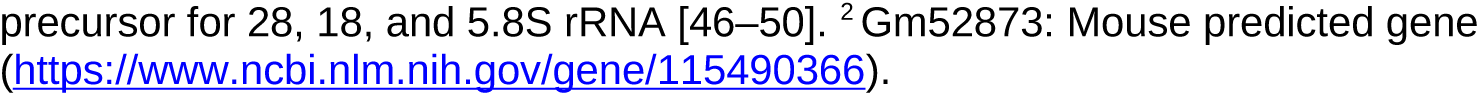
Candidate circRNA regions identified as commonly down-regulated by tamoxifen and letrozole exposure in mouse mammary gland.

**Table 3.** Mouse database circRNA identifier listings for candidate differentially expressed circRNAs up-regulated by both tamoxifen and letrozole in mouse mammary gland.

|  | Host Gene(s) in region of<br>Identified circRNA | Circ-Atlas | Circ-Base | Circ-<br>bank | CIRC-pedia | TSCD |
| --- | --- | --- | --- | --- | --- | --- |
| 1 | ***Fmn1*** | mmu-Fmn1_0001,<br>mmu-Fmn1_0006,<br>mmu-Fmn1_0040 |  | mmu_Fmn1_0032000,<br>mmu_Fmn1_0034000,<br>mmu_Fmn1_0044000 |  |  |
| 2 | ***Gm34744, Bcam*** | mmu-Bcam_0001 |  | mmu_Bcam_0009000 | CIRCMMU_Bcam_3 | heart_14wks_SRR1772419 |
| 3 | ***Hmg20a*** | mmu-Hmg20a_0007 |  | mmu_Hmg20a_0006000 | CIRCMMU_Hmg20a_1 |  |
| 4 | ***Scn4a*** |  |  |  |  |  |

**Table 4.**
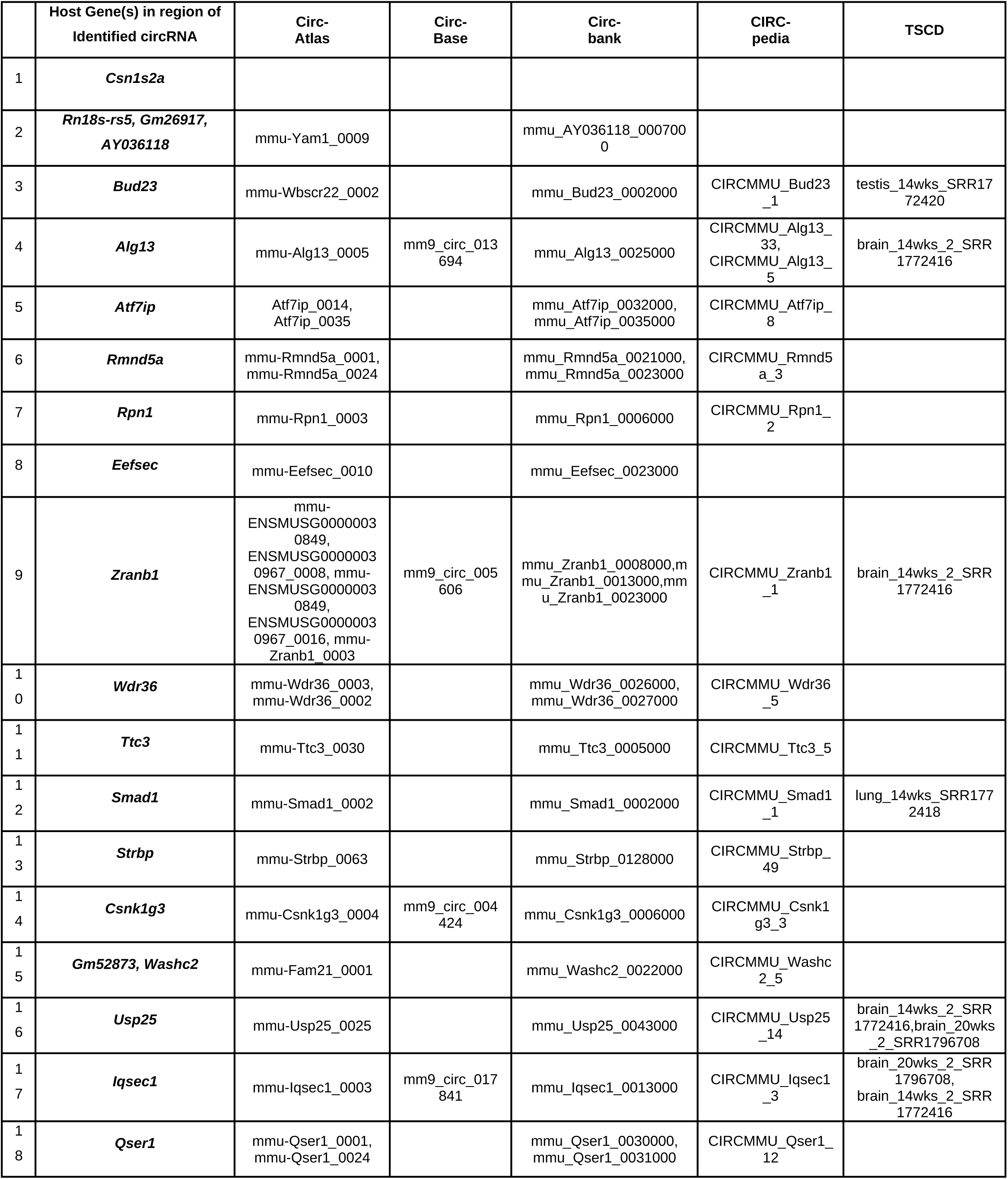
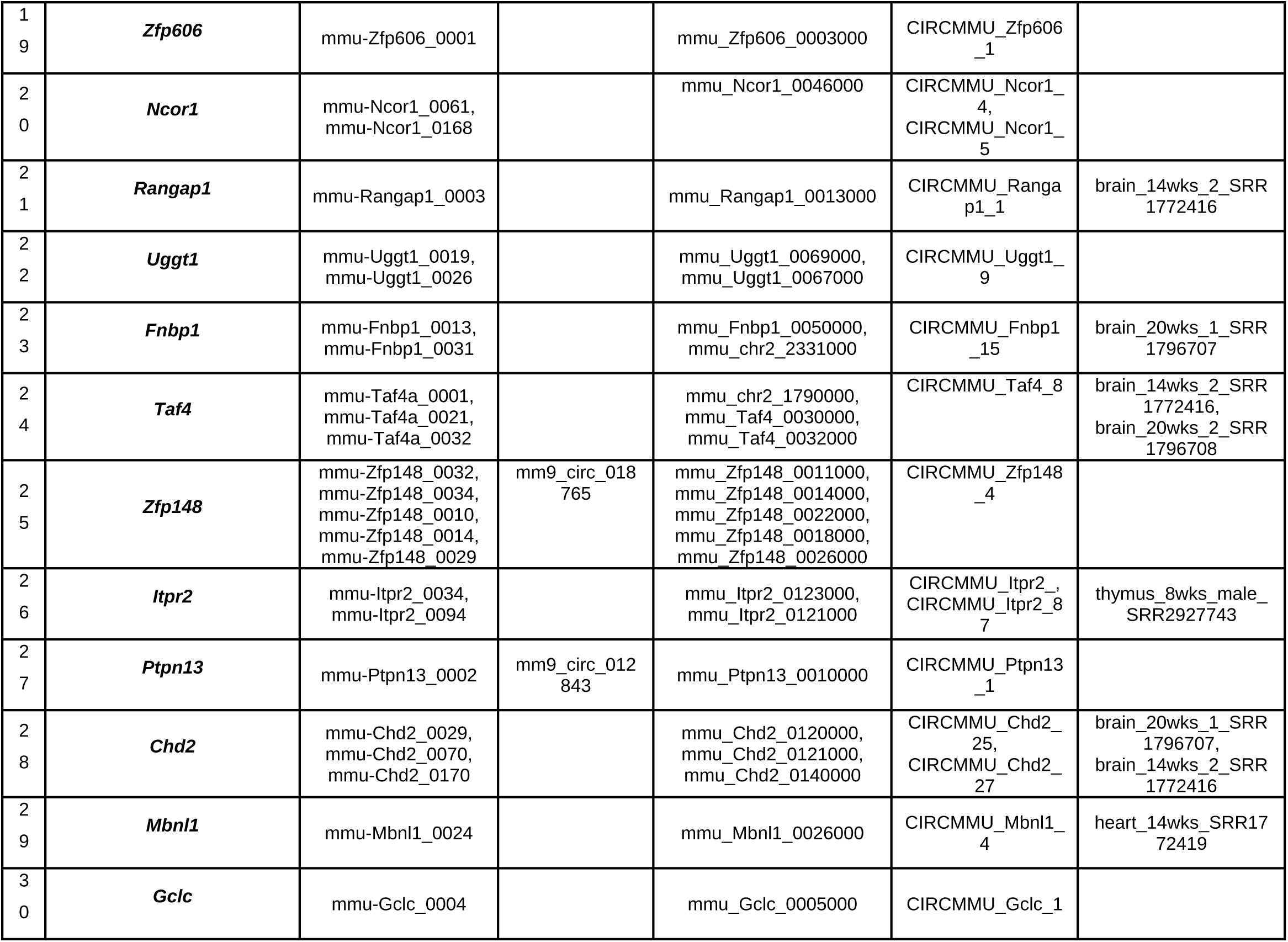
Mouse database circRNA identifier listings for candidate differentially expressed circRNAs down-regulated by both tamoxifen and letrozole in mouse mammary gland.

**Table 5.** Candidate circRNAs commonly up-regulated by tamoxifen and letrozole exposure are derived from host genes known to be expressed in human breast epithelial cells. Human database circRNA identifier listings presented for orthologous regions.

|  | Host Gene | Human Host Gene Name | Host Gene |  | Human circRNA orthologous to identified mouse circRNA |  |  |  |  |
| --- | --- | --- | --- | --- | --- | --- | --- | --- | --- |
|  |  |  | Host gene expressed in human breast tissue* | Host gene expression in primary human breast epithelial cells (avgFPKM) (67) | circAtlas [mean expression (counts per million) shown for human breast if available] | circBase | Circbank | CIRCpedia | TSCD |
| 1 | **Fmn1** | Formin 1 | yes | 1.44 | hsa-FMN1_0002 [0.036], hsa-FMN1_0024, hsa-FMN1_0037, hsa-FMN1_0071 |  | hsa_FMN1_0012400, hsa_FMN1_0012500, hsa_FMN1_0012600, hsa_FMN1_0012100 |  |  |
| 2 | **Gm34744, Bcam** | Basal cell adhesion molecule (Lutheran blood group) | Gm34744: no (mouse specific), Bcam: yes | Gm34744: no (mouse specific), Bcam: 166.24 |  |  |  |  |  |
| 3 | **Hmg20a** | High mobility group 20A | yes | 12.29 | hsa-HMG20A_0003 | hsa_circ_0036428 | hsa_HMG20A_0001200 | CIRCHSA_HMG20A_11 |  |
| 4 | **Scn4a** | Sodium voltage-gated channel alpha subunit 4 | yes | Not detected. |  |  |  |  |  |

\*https://www.proteinatlas.org/. Abbreviations: avgFPKM: (Average Fragments Per Kilobase of transcript per Million mapped reads) for n=18 independent samples of primary human high-risk breast cells)<sup>51</sup>.

**Table 6.**
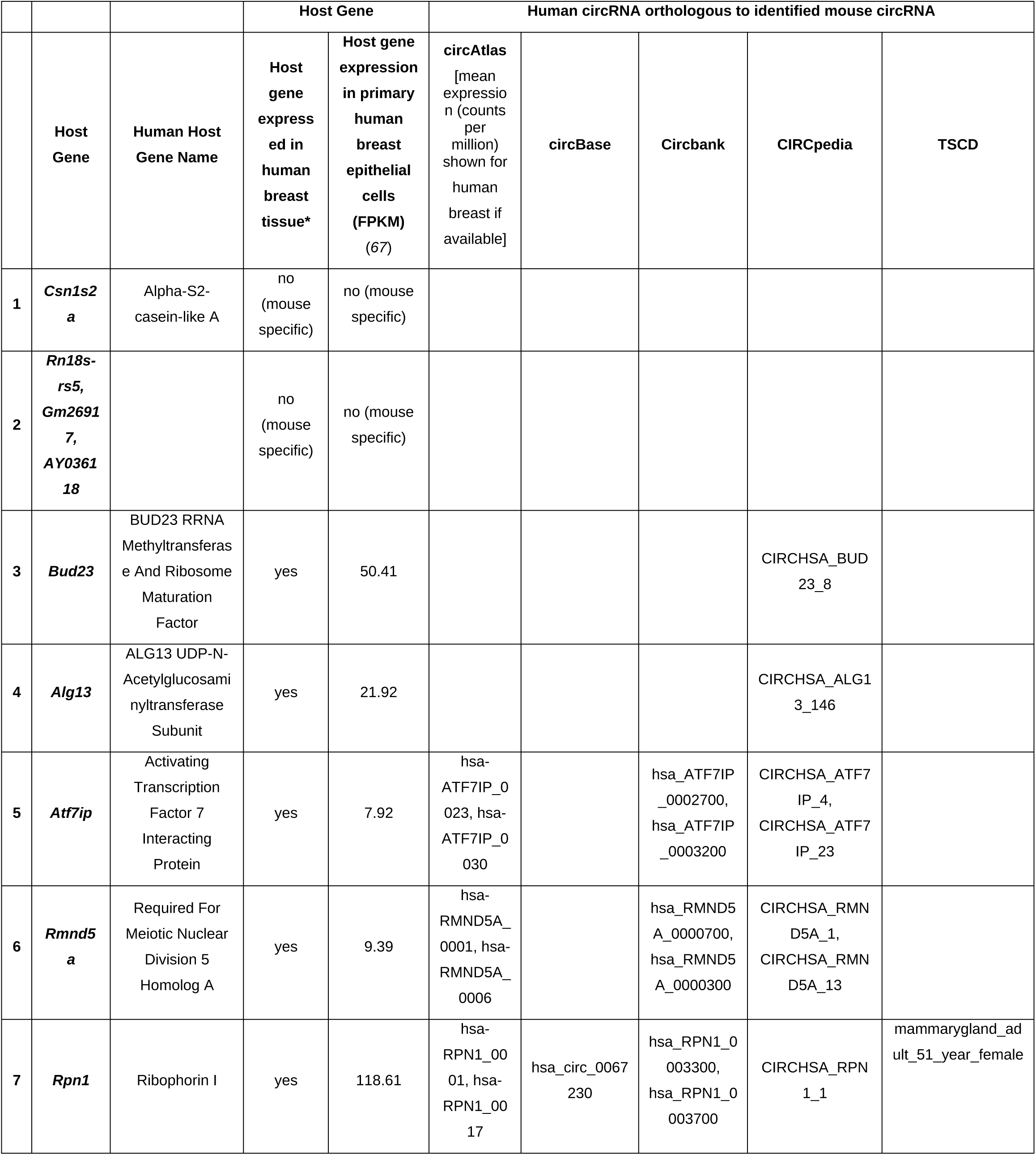
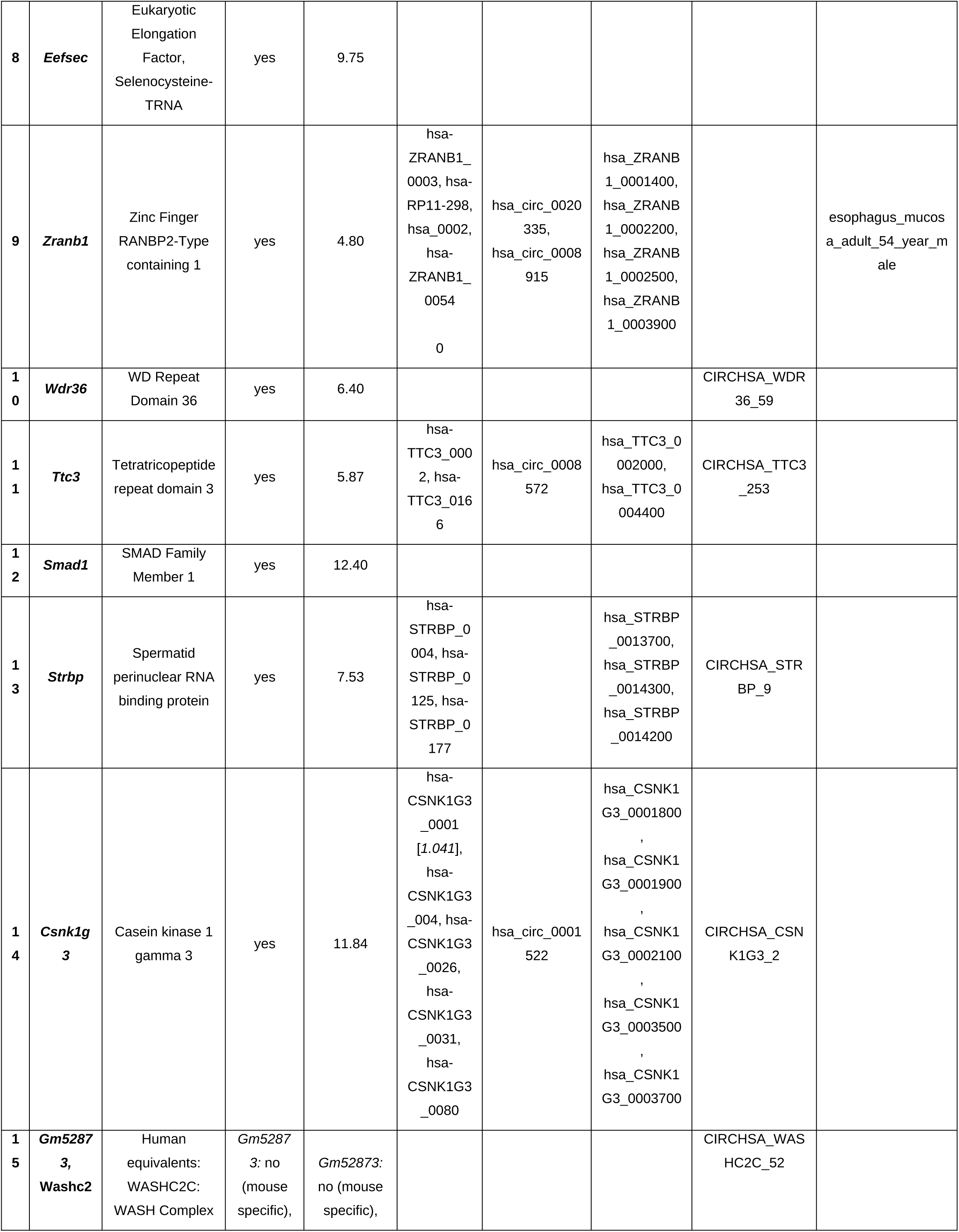
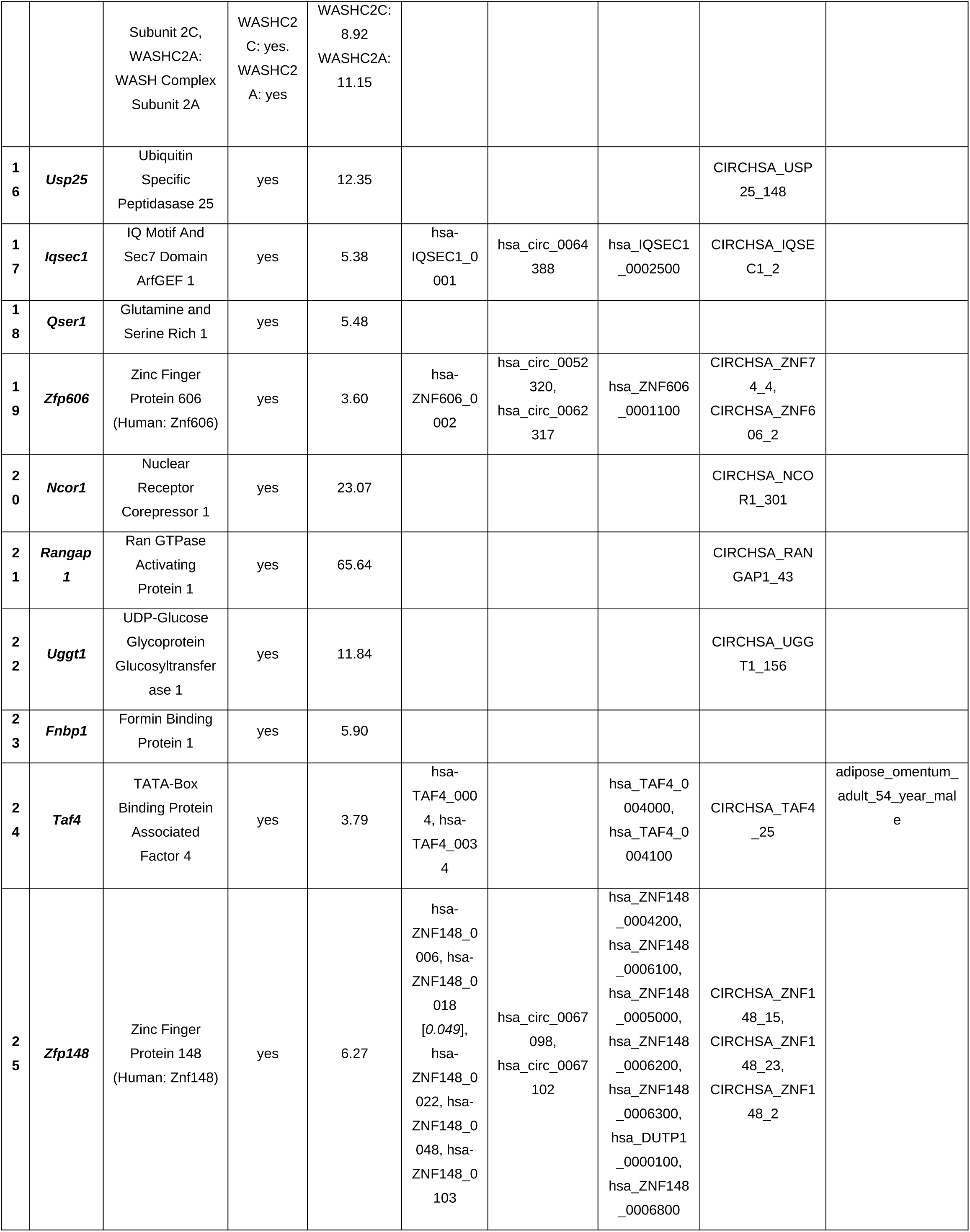
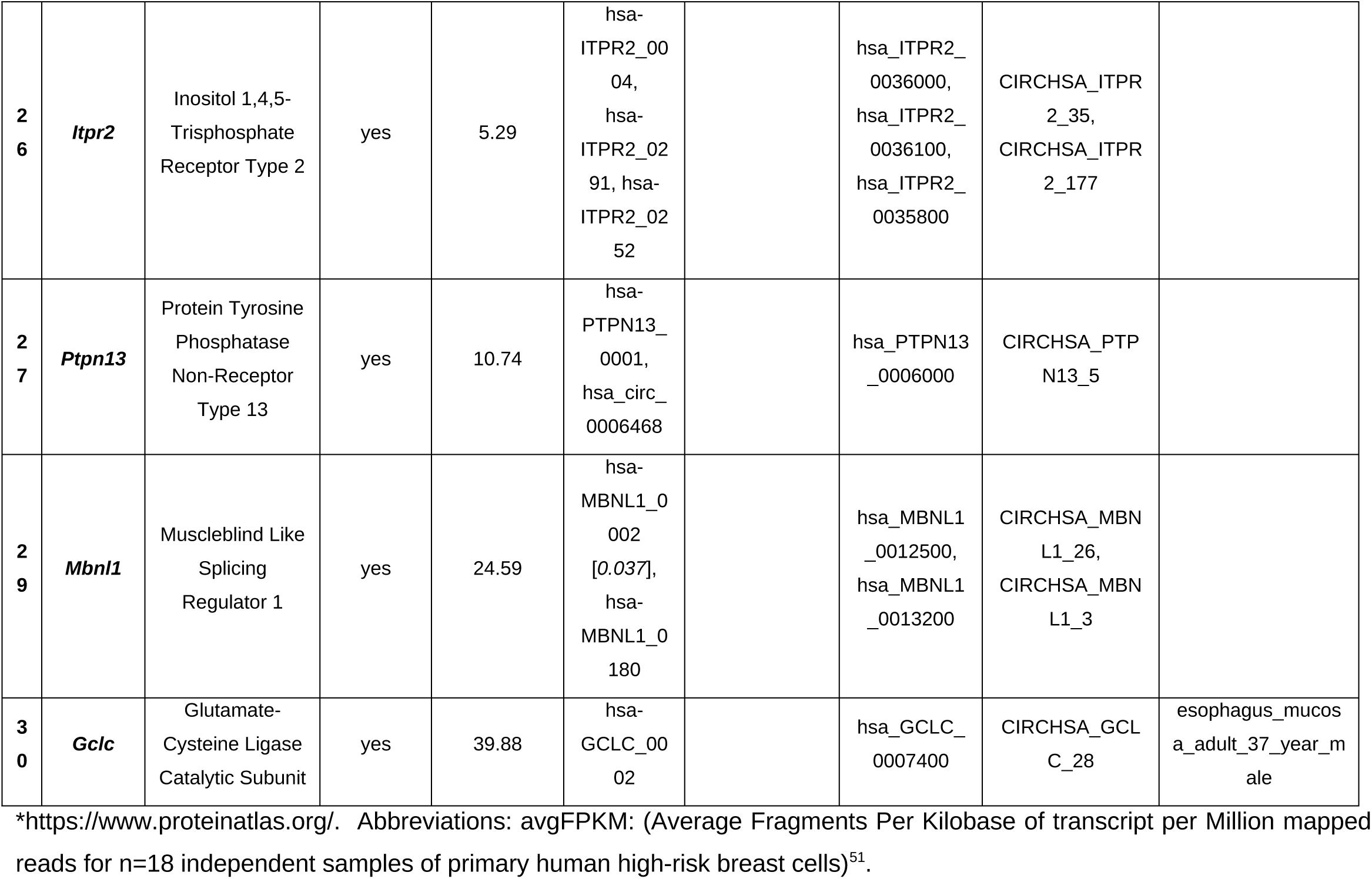
Candidate circRNAs commonly down-regulated by tamoxifen and letrozole exposure are derived from host genes known to be expressed in human breast epithelial cells. Human database circRNA identifier listings presented for orthologous regions.

## Discussion

Using a bioinformatic approach, this study demonstrated that there is a set of circRNAs expressed in mammary tissue that undergo statistically significant changes in expression coincident with a positive therapeutic response to antihormonals. Development of a circRNA panel that could serve as biomarkers for a positive therapeutic response could help identify early treatment failures for current agents and assist in validating new anti-hormonals currently under development (*31*). Currently, clinical strategies for assessing an early therapeutic response to anti-hormonals rely on measuring estrogen levels following aromatase inhibitor treatment to ensure they drop (*32*) or determining if breast density decreases with anti-hormonal exposure (*33*, *34*). Because changes in circRNA expression can be measured in circulating blood (*35–37*), they represent a non-invasive means of assessing the molecular response to anti-hormonals.

Preclinical development programs in oncology commonly exploit translational models for initial identification and characterization of circRNA biomarkers (*38–40*). Here, two GEMMs were used to increase generalizability of the bioinformatic analyses by validating if the same circRNAs were differentially expressed across two different mechanisms of human breast cancer generation and two different classes of anti-hormonal agents. The design of the bioinformatic pipeline for identification of candidate circRNAs was optimized for rigor and reproducibility by focusing only on circRNAs found to be present in all biological samples tested and filtering for circRNAs that exhibited at least a 2-fold difference in expression levels. Current recommendations suggest that any biomarker test utilizing circRNAs include at least three distinct circRNAs (*10*, *11*, *41*). Here, we identified 25 distinct candidate orthologous circRNAs that are derived from host genes expressed in human breast, a reasonable collection to begin paring down to find at least three circRNAs that might exhibit measurable changes in expression in human peripheral blood. These experimental results show a proof of principle supporting direct studies in human tissue to discover even more circRNAs relevant to hormone response in both breast and other reproductive tissues, including the possibility of using circRNAs as therapeutic agents (*37*).

Limitations of the study include the relatively small number of each genotype/condition set of samples (n=3). In this study the small sample size was mitigated by combining together both GEMM for identification of statistically significant changes in circRNA expressions, increasing the sample size for each comparison to n=6. The study was designed as a bioinformatic analysis to establish proof of principle. Future studies will be needed to validate expression changes directly in mouse and human tissue. In addition, while this study focused on differential circRNA abundance, we did not test whether the identified candidates engage in miRNA sponging or other miRNA-linked mechanisms, which will require dedicated computational prediction and experimental validation (*42–44*). For a circRNA to be an optimal biomarker, it should be secreted in an exosome so that it may be trafficked to the peripheral blood. The candidate circRNAs identified in this study have not yet been assessed for their exosome secretion patterns. Together, these results position circRNAs as a promising, mechanism-agnostic molecular readout of effective estrogen-pathway suppression, establishing a rational foundation for future translational validation of a minimally invasive biomarker panel.

## Methods

### Mouse Models and Total RNA Sequencing Data

Total single-ended RNA sequencing data analyzed in this project was downloaded from the National Center for Biotechnology Information’s Gene Expression Omnibus (GSE70440, GSE201767). Total RNA sequencing fastq files were obtained from mammary tissue obtained from cohorts of mammary tumor virus–reverse tetracycline–controlled transactivator/Tet-operator (*tet-op*)–*Esr1* and mouse mammary tumor virus–reverse tetracycline–controlled transactivator/*tet-op*–*CYP19A1* mice on a C57Bl/6 background from experiments examining the impact of age, *Esr1* or *CYP19A1* over-expression, or exposure to tamoxifen or letrozole (n=3 independent mammary gland samples per condition) (*15*, *25*). Studies on the mice were approved by the Georgetown University Animal Care and Use Committee (Protocol number 2018-0022, Progression and regression of neoplasia in aged mice, protocol approval date: 05/01/2018).

### Identification of circRNAs differentially regulated by exposure to either tamoxifen or letrozole using the nf-core/circRNA pipeline

To identify circRNAs differentially regulated by tamoxifen exposure in both models, an nf-core/circrna detection pipeline (*13*, *14*) was used on the total single-ended RNA sequencing data from mammary glands of tet-op-Esr1 and tet-op-CYP19A1 mouse models (age 20 months) with and without a two-month exposure to tamoxifen, combined data from the mammary glands of tet-op-Esr1 and tet-op-CYP19A1 mouse models (age 20 months) with and without a two-month exposure to letrozole, combined data from mammary glands of tet-op-Esr1 mice with and without induction of the tet-op-Esr1 transgene using GRCm39 as the reference genome (https://www.ncbi.nlm.nih.gov/datasets/genome/GCF_000001635.27/, accessed 11/20/2025). Sequencing was performed using the Illumina NextSeq 550, SE 75-bp read length; minimum reads 50 million per sample. The nf-core/circrna detection pipeline utilizes different tools for back-splice junction (BSJ) detection (CIRIquant (*45*), CircExplorer2 (*46*), CircRNA finder (*47*), DCC (*48*), Find circ (*49*), MapSplice (*50*), Segemehl (*51*)), annotates the detected circRNAs using Gene Transfer Format (GTF)-based and database-based annotation and extracts the sequences of the circRNAs and quantifies their expression. For database annotation, circBase (*52*) and circAtlas (*53*) were used. circBase was lifted over from its original reference to the mm39 reference genome using the University of California Santa Cruz (UCSC) liftOver tool (*54*). The length of the overlap (latest start and the first end of the identified differentially expressed circRNA and the database entry) has to be at least 90% of the length to be considered a match.

### Differential expression analysis and identification of reproducibly differentially expressed circRNAs under different hormonal conditions

DESeq2 (*55*) was used to identify differentially expressed circRNAs (*56–58*) in two analyses. The pipeline detected 56,621 unique circRNAs (defined by chr:start–stop:strand coordinates). Due to the sparsity of the raw count data, circRNAs were pre-filtered to retain only those with at least 10 counts in at least three samples, resulting in 13,803 candidates for differential expression testing. For both analyses, the design formula “∼ age + transgene + induction + drug” was used to control for potential confounding effects of age, transgene status, and induction while assessing drug-associated expression changes. Differential expression was evaluated using contrasts comparing treated versus untreated samples (“contrast = c(“drug”, “tamoxifen”, “no”)” and “contrast = c(“drug”, “letrozole”, “no”)”). Log2 fold change shrinkage (*55*) was applied to account for the high variance in effect size estimation, and circRNAs with an absolute log2 fold change greater than 1 were considered for interpretation. P-values were adjusted for multiple testing using the Benjamini–Hochberg method (*59*), and circRNAs with an adjusted p-value (padj) < 0.05 were considered statistically significant. To find circRNAs that were reproducibly differentially regulated after both tamoxifen or letrozole exposure, circRNAs that were significantly differentially regulated in the same direction with both tamoxifen and letrozole exposure were identified. Chromosomal coordinates had to be within 1 basepair to be considered the same circRNA, and detection of the BSJ had to be found in both + and – amplified cDNA strands of the RNAseq.

### Database Annotation

Initial database annotation was performed as part of the nf-core/circrna detection pipeline. Subsequently, a secondary database annotation was performed on all candidate circRNAs identified. These circRNA entries were extracted and stored in BED format. Databases used for the secondary annotation were circAtlas 3.0 (https://ngdc.cncb.ac.cn/circatlas/, accessed 23 November 2025) (*53*), CircBase (https://www.circbase.org/, accessed 23 November 2025 (*52*); Circbank (https://www.circbank.cn/#/home, accessed 23 November 2025) (*60*), CIRCpedia v3 (https://bits.fudan.edu.cn/circpediav3/, accessed 23 November 2025) (*61*), and Tissue-Specific CircRNA Database (TSCD) (http://gb.whu.edu.cn/TSCD/, accessed 23 November 2025) (*62*). To conduct the secondary annotation, all available mouse and human circRNA annotations were downloaded from the circAtlas 3.0 (*53*), CircBase (*52*), CIRCpedia v3 (*61*), and TSCD databases. The downloaded database files were then converted into a standard BED format for subsequent analyses. Genomic coordinates from the database files were mapped to the target assembly using the UCSC LiftOver tool, which converts genomic coordinates between assemblies and identifies corresponding aligned regions (*54*). Human coordinates from the hg19 and hg38 genome assemblies were lifted to the mm39 mouse genome assembly, and mouse entries were lifted from mm9 and mm10 to mm39, ensuring all comparisons used the most current mouse reference genome. For cross-species liftovers from human to mouse, the minMatch LiftOver parameter was reduced to 0.1 to maximize retrieval of putative orthologous regions. All default parameters were used for mouse genome conversions. It should be noted that the LiftOver tool identifies approximate homologous genomic regions, but does not assess sequence similarity or functional equivalence. Consequently, the resulting mappings do not account for sequence conservation, nor do they imply functional orthology. Once all of the database files were converted to a compatible reference assembly, overlaps between the candidate circRNA data and database annotations were assessed using bedtools intersect (*63*). Bedtools intersect identifies overlapping genomic features and provides flexible parameters for defining and reporting these overlaps. The parameters -f 0.9 -r -loj -wa -wb were used, ensuring that circRNAs were only considered a match if their genomic coordinates overlapped by at least 90%. The same parameters were applied to all human and mouse datasets.

### Characterization of host gene expression in mouse mammary gland and human breast

To evaluate host gene expression levels for the identified circRNAs, RNAseq data was analyzed to determine transcripts per million (TPM) values in mammary glands of 20-month-old female mice without exposure to tamoxifen or letrozole (n=3 each genotype) for each host gene (GSE70440) (*25*). Mean TPM values for each genotype/condition were calculated using https://www.calculatorsoup.com/calculators/statistics/mean-median-mode.php. In an adaptation of a previously published method (*64*), relative circular over linear ratios (CLR) were calculated (circular basemean/linear TPM) using the basemean levels calculated by the pipeline for the coordinates of the identified circRNAs over transcripts per million (TPM) of the linear forms, a value that corrected for both sequencing depth (total read count) and gene length for the linear forms. Evaluation of host gene differential expression following tamoxifen or letrozole exposure was determined using DEseq2 as described previously (n=3 each genotype/condition) (*25*). For evaluation of host gene expression in human breast tissue, breast tissue specific RNA expression data from The Human Protein Atlas https://www.proteinatlas.org (last accessed February 6, 2026) was recorded (*65*, *66*) and average fragments per kilobase of transcript per million mapped reads (avgFPKM) for n=18 independent samples of primary human high-risk breast cells determined as described previously (*67*). Gene Set Enrichment Analysis (GSEA) was performed on mouse and human sets of host genes for the circRNAs identified for assessment of statistically significant relationships with known gene sets (hallmark gene sets, C3 regulatory target gene sets, C2 curated gene sets, M3 regulatory target gene sets) https://www.gsea-msigdb.org/gsea/msigdb/index.jsp (accessed February 5, 2026) (*28*, *68–70*).

Nico: Data curation, Formal analysis, Methodology, Software

Malte: Data curation, Formal analysis, Investigation, Methodology, Software, Validation, Visualization, Writing – review & editing

PR: Investigation, Methodology, Writing – original draft, Writing – review & editing

PAF: Conceptualization, Funding acquisition, Investigation, Methodology, Resources, Supervision, Validation, Writing – original draft, Writing – review & editing

MH: Conceptualization, Funding acquisition, Investigation, Methodology, Project administration, Software, Supervision, Writing – original draft, Writing – review & editing

ML: Conceptualization, Funding acquisition, Investigation, Methodology, Project administration, Resources, Software, Supervision, Validation, Writing – review & editing

---

**Supplementary Table 1.** Expression patterns of host genes in mouse mammary gland for candidate circRNAs identified as commonly up-regulated by tamoxifen and letrozole exposure.

|  | Host Gene | Baseline Mean TPM | Relative Circular-to-Linear Ratio (CLR) | Significantly differentially regulated with letrozole exposure in *CYP19A1* mice | Significantly differentially regulated with letrozole exposure in *Esr1* mice | Significantly differentially regulated with tamoxifen exposure in *CYP19A1* mice | Significantly differentially regulated with tamoxifen exposure in *Esr1* mice |
| --- | --- | --- | --- | --- | --- | --- | --- |
| 1 | ***Fmn1*** | 3.89 | 0.12 | no | Sig. down-regulated (log2fold -0.56) | no | no |
| 2 | ***Gm34744*<sup>1</sup>,**<br>***Bcam*** | *Bcam*: 29.69<sup>2</sup> | 0.04<sup>3</sup> | *Bcam*: no | *Bcam*: Sig. up-regulated (log2fold 0.51) | *Bcam*: no | *Bcam*: Sig. up-regulated (log2fold 0.53) |
| 3 | ***Hmg20a*** | 11.27 | 0.14 | no | no | no | no |
| 4 | ***Scn4a*** | 28.09<sup>4</sup> | 0.04 | no | no | Sig. down-regulated (log2fold -0.50) | Sig. up-regulated (log2fold 1.23) |

Data analyzed from dataset GSE201326 (15). Differentially expressed genes (DEGs) between mice exposed to either letrozole or tamoxifen versus controls identified using DESeq2. Genes were considered statistically significantly differentially expressed when the adjusted P value was <0.05. <sup>1</sup>*Gm34744*: full length gene expression not detected in dataset GSE201326. <sup>2</sup> Expressed statistically significantly higher (log2fold 0.61) in *CYP19A1* (mean 35.57) vs. *Esr1* (mean 23.81) mice. <sup>3</sup> Calculated using only *Bcam* TPM as linear *Gm34744* not identified in the analysis of the RNAseq data for linear forms. <sup>4</sup> Expressed statistically significantly higher (log2fold 1.10) in *CYP19A1* (mean 37.72) vs. *Esr1* (mean 18.46) mice. Abbreviations: *Esr1*: *MMTV-rtTA/Tet-op-Esr1*. *CYP19A1*: *MMTV-rtTA/Tet-op-CYP19A1*. Baseline Mean TPM: Mean TPM (Transcripts per million) in mammary gland tissue from 20-month-old female mice without exposure to tamoxifen or letrozole (n=6 samples).

**Supplementary Table 2.** Expression patterns of host genes in mouse mammary gland for candidate circRNAs identified as commonly down-regulated by tamoxifen and letrozole exposure.

|  | Host Gene | Baseline Mean TPM | Relative Circular-to-Linear Ratio (CLR) | Significantly differentially regulated with letrozole exposure in *CYP19A1* mice | Significantly differentially regulated with letrozole exposure in *Esr1* mice | Significantly differentially regulated with tamoxifen exposure in *CYP19A1* mice | Significantly differentially regulated with tamoxifen exposure in *Esr1* mice |
| --- | --- | --- | --- | --- | --- | --- | --- |
| 1 | ***Csn1s2a*** | 3790.42 | 0.06 | no | Significantly down-regulated (log2fold -9.22) | no | no |
| 2 | ***Rn18s-rs5*,**<br>***Gm26917*,**<br>***AY036118*** | 1062.89 | 0.01 | no | no | no | no |
| 3 | ***Bud23*** | 13.93 | 0.33 | no | no | no | no |
| 4 | ***Alg13*** | 5.02 | 0.60 | no | no | no | no |
| 5 | ***Atf7ip*** | 29.04<sup>1</sup> | 0.08 | no | no | no | no |
| 6 | ***Rmnd5a*** | 32.54<sup>2</sup> | 0.08 | no | Significantly up-regulated (log2fold 0.33) | no | no |
| 7 | ***Rpn1*** | 49.78<sup>3</sup> | 0.05 | no | Significantly down-regulated (log2fold -0.59) | no | Significantly down-regulated (log2fold -0.63) |
| 8 | ***Eefsec*** | 4.98 | 0.43 | no | no | no | no |
| 9 | ***Zranb1*** | 17.65 | 0.12 | no | no | no | no |
| 10 | ***Wdr36*** | 12.39 | 0.19 | no | no | no | no |
| 11 | ***Ttc3*** | 30.87 | 0.07 | no | no | no | no |
| 12 | ***Smad1*** | 9.03 | 0.34 | no | no | no | no |
| 13 | ***Strbp*** | 10.78<sup>4</sup> | 0.15 | no | Significantly down-regulated (log2fold -0.76) | no | Significantly down-regulated (log2fold -0.55) |
| 14 | ***Csnk1g3*** | 6.66 | 0.48 | no | no | no | no |
| 15 | ***Gm52873*,**<br>***Washc2*<sup>5</sup>** | *Washc2*: 25.02 | *Washc2*: 0.09 | *Washc2*: no | *Washc2*: no | *Washc2*: no | *Washc2*: no |
| 16 | ***Usp25*** | 29.90 | 0.05 | no | Significantly up-regulated (log2fold | no | no |
|  |  |  |  |  | 0.35) |  |  |
| 17 | ***lqsec1*** | 19.98<sup>6</sup> | 0.09 | no | Significantly up-regulated (log2fold 0.71) | no | Significantly up-regulated (log2fold 0.74) |
| 18 | ***Qser1*** | 9.66 | 0.23 | no | no | Significantly down-regulated (log2fold -0.43) | no |
| 19 | ***Zfp606*** | 6.17 | 0.31 | no | no | no | no |
| 20 | ***Ncor1*** | 84.93 | 0.02 | no | no | no | no |
| 21 | ***Rangap1*** | 10.82 | 0.22 | no | no | no | no |
| 22 | ***Uggt1*** | 24.15 | 0.09 | no | no | no | no |
| 23 | ***Fnbp1*** | 17.28 | 0.09 | no | no | no | no |
| 24 | ***Taf4*** | 5.44 | 0.28 | no | no | no | no |
| 25 | ***Zfp148*** | 16.52 | 0.06 | no | no | no | no |
| 26 | ***Itpr2*** | 15.76 | 0.14 | no | no | no | no |
| 27 | ***Ptpn13*** | 4.63 | 0.34 | no | no | no | no |
| 28 | ***Chd2*** | 41.98 | 0.002 | no | no | no | no |
| 29 | ***Mbnl1*** | 126.26 | 0.008 | no | Significantly up-regulated (log2fold 0.51) | no | no |
| 30 | ***Gclc*** | 12.10 | 0.19 | no | no | Significantly down-regulated (log2fold -0.72) | no |

Data analyzed from dataset GSE201326 [4]. Differentially expressed genes (DEGs) between mice exposed to either letrozole or tamoxifen versus controls identified using DESeq2. Genes were considered statistically significantly differentially expressed when the adjusted P value was <0.05. <sup>1</sup> Expressed statistically significantly lower (log2fold -0.50) in *CYP19A1* (mean 22.68) vs. *Esr1* (mean 35.39) mice. <sup>2</sup> Expressed statistically significantly higher (log2fold 0.40) in *CYP19A1* (mean 36.22) vs. *Esr1* (mean 28.85) mice. <sup>3</sup> Expressed statistically significantly lower (log2fold -0.37) in *CYP19A1* (mean 43.06) vs. *Esr1* (mean 56.48) mice. <sup>4</sup> Expressed statistically significantly lower (log2fold -0.41) in *CYP19A1* (mean 8.71) vs. *Esr1* (mean 12.85) mice. <sup>5</sup> Only linear *Washc2* detected in dataset GSE201326. <sup>6</sup> Expressed statistically significantly higher (log2fold 0.77) in *CYP19A1* (mean 42.51) vs. *Esr1* (mean 15.45) mice. Abbreviations: *Esr1*: MMTV-rtTA/Tet-op-*Esr1*. *CYP19A1*: MMTV-rtTA/Tet-op-*CYP19A1*.

